# Synchronous spiking of cerebellar Purkinje cells during control of movements

**DOI:** 10.1101/2021.09.16.460700

**Authors:** Ehsan Sedaghat-Nejad, Jay S. Pi, Paul Hage, Mohammad Amin Fakharian, Reza Shadmehr

**Affiliations:** Laboratory for Computational Motor Control, Dept. of Biomedical Engineering, Johns Hopkins School of Medicine, Baltimore, Maryland; School of Cognitive Sciences, Institute for Research in Fundamental Sciences, Tehran, Iran

## Abstract

The information that the brain transmits from one region to another is often viewed through the lens of firing rates. However, if the output neurons could vary the timing of their spikes with respect to each other, then through synchronization they could highlight information that may be critical for control of behavior. In the cerebellum, the computations that are performed by the cerebellar cortex are conveyed to the nuclei via inhibition. Yet, synchronous activity entrains nucleus neurons, making them fire. Does the cerebellar cortex rely on spike synchrony within populations of Purkinje cells (P-cells) to convey information to the nucleus? We recorded from multiple P-cells while marmosets performed saccadic eye movements and organized them into populations that shared a complex spike response to error. Before movement onset, P-cells transmitted information via a rate code: the simple spike firing rates predicted the direction and velocity of the impending saccade. However, during the saccade, the spikes became temporally aligned within the population, signaling when to stop the movement. Thus, the cerebellar cortex relies on spike synchronization within a population of P-cells, not individual firing rates, to convey to the nucleus when to stop a movement.

## Introduction

To understand how neurons in a region of the brain respond to sensory information or participate in control of movements, we typically search for correlates of the sensory and motor variables in the patterns of spikes. These patterns are usually quantified via the average firing rates of neurons. However, there can be additional information in the timing of each spike, as exemplified by the independent rate and temporal codes in the hippocampus ^1^, the thalamus ^2^, and the somatosensory cortex ^3^. A central question is whether neurons use spike timing to transmit functionally relevant information from one region of the brain to another.

A special form of temporal coding is synchronization of spikes among a group of neurons. For example, synchronization among glutamatergic thalamic neurons increases the efficiency of driving post-synaptic neurons in the somatosensory cortex ^4^. However, unlike the thalamus, the sole output from the cerebellar cortex is via GABAergic Purkinje cells (P-cells). As a result, asynchronous activity of P-cells inhibits the cerebellar nucleus neurons. Indeed, previous analysis of spike timing in single P-cells did not find evidence that timing of spikes affected ongoing movements ^5^. Yet, there are specialized mechanisms in the cerebellar cortex that promote synchronization of nearby P-cells ^6^, raising the question of whether the cerebellum relies on synchronization to transfer information from its cortex to its nuclei.

In principle, when a population of P-cells synchronizes their spikes, they can drive cerebellar output in a way that is not possible via asynchronous spiking ^7^. For example, when P-cells are synchronously stimulated (in slice, and anesthetized preparations), they entrain the nucleus cells, transforming their inhibitory inputs to the nucleus into production of spikes ^8,9^. This raises the possibility that analogous to the thalamic input to the cerebral cortex, P-cells may rely on synchronization to convey information to the nucleus, possibly affecting a specific part of the ongoing movement ^10^.

Here, we focused on saccadic eye movements because they are so brief as to preclude the possibility of sensory feedback, requiring the brain to rely entirely on its internal predictions ^11–13^. These predictions depend critically on the cerebellum ^14,15^. For example, firing rates of populations of P-cells, but not individual cells, predict the direction and velocity of the ongoing saccade ^16,17^. However, to check for synchrony we needed to simultaneously record from multiple P-cells during saccades, something that to our knowledge had not been accomplished in any primate species.

## Results

We focused on marmosets, a primate that like macaques and humans relies on saccadic eye movements to explore its visual scene, but is a fraction of the size of macaques, thus making it possible to record from the cerebellum using short, multi-channel probes. We used MRI and CT-aligned maps of each animal’s cerebellum ^18^ to guide electrodes and record from P-cells in lobule VI and VII of the vermis (Fig. 1C). Because there were no previous electrophysiological data from the marmoset cerebellum, we searched for saccade related activity and found that P-cells in the posterior lobule VI and anterior lobule VII produced simple spikes that were modulated during saccades. Thus, we focused on these regions and recorded from n=149 well-isolated P-cells (Supplementary Fig. S1 provides characteristics of the entire data set). Crucially, our data included n=42 pairs of simultaneously isolated P-cells that were recorded from separate channels.

**Fig. 1.**
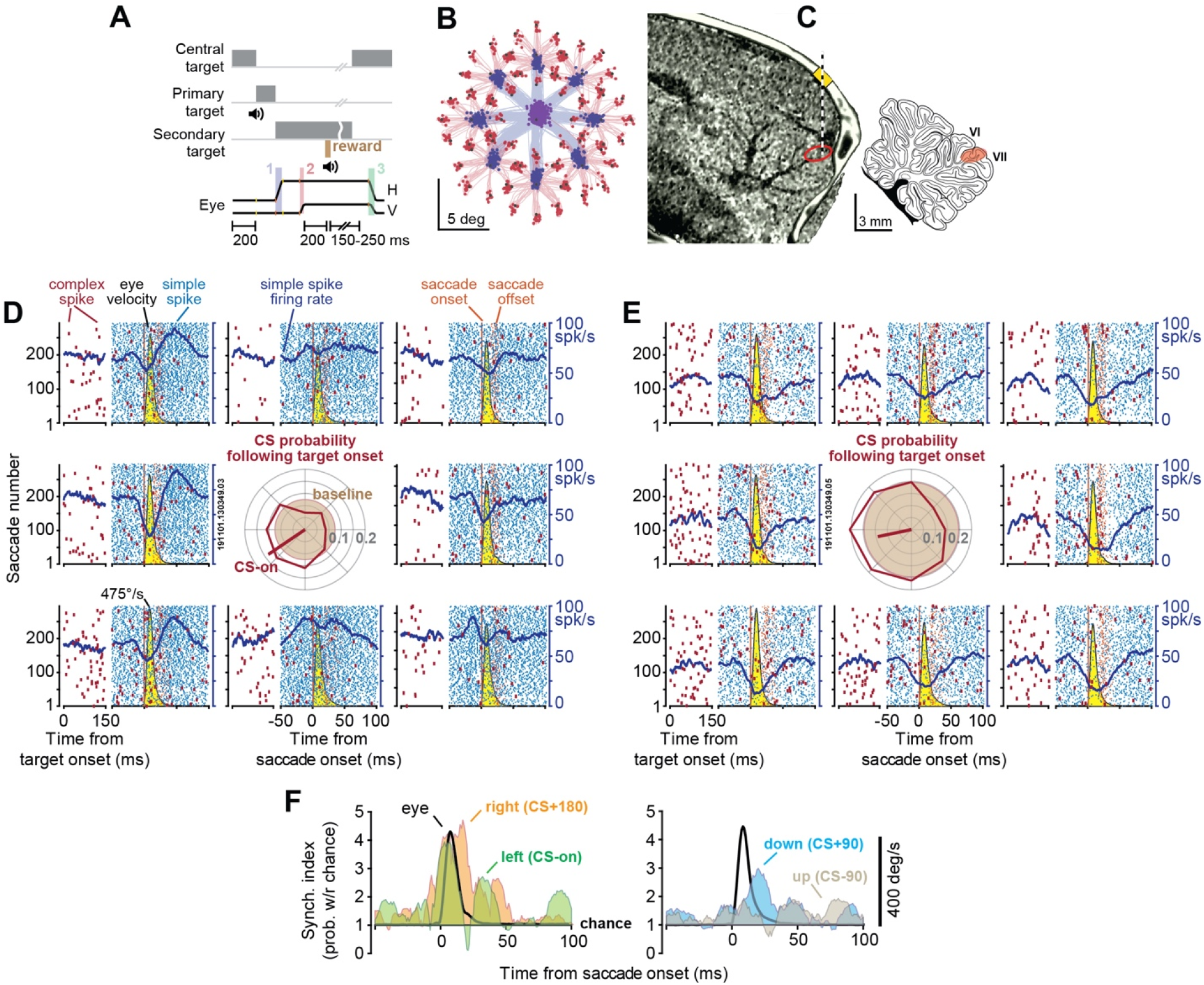
P-cells synchronized their simple spikes during saccades. **A**. Experimental paradigm. Marmosets were trained to make saccades to visual targets that appeared randomly at one of 8 directions. Onset of the primary saccade (labeled 1 in the lowest trace) resulted in the replacement of the primary target with a secondary target, also at a random direction. Following the secondary saccade (labeled 2), and a fixation period, reward was delivered, and the center target was displayed, resulting in a centripetal saccade (labeled 3). **B**. Eye position for the primary (blue) and secondary (red) saccades in a typical experiment. **C**. We used the MRI and CT images of each animal to guide the electrodes to lobule VI or VII of the cerebellar vermis. **D** & **E**. Simple (blue) and complex spikes (red) in two simultaneously recorded P-cells during saccades to various directions. Eye velocity is shown via the yellow curve. The complex spikes are also aligned to the onset of the visual target. Both cells exhibited a reduction in simple spikes during saccades, with a modulation pattern that lasted much longer than the saccade. CS probability during the 200ms following target onset is quantified via the center plot. Baseline CS probability is shown by the brown circle at center. The target direction that produces the highest CS probability (CS-on) is estimated by the red line at center. **F**. Synchronization index during saccades to various directions. This index quantified the probability of synchronization with respect to chance at 1 ms time bins. Eye velocity is indicated by the black curve. Probability of synchronization is greatest for saccades in direction CS+180, reaching a peak at around saccade deceleration.

We trained the animals to fixate a central target and make a saccade to a primary target that appeared at random in one of 8 directions (Fig. 1A & 1B). At the onset of the primary saccade the target was erased and replaced with a secondary target, also at a random location. Following a random period of fixation and delivery of reward, the secondary target was erased, the central target was displayed, and the trial re-started. Whereas production of saccades accompanied modulation of simple spikes (SS), the random nature of the visual stimuli produced sensory prediction errors, promoting modulation of complex spikes (CS) ^16,17,19,20^.

Data from a pair of simultaneously recorded P-cells are shown in Figs. 1D & 1E. Despite their proximity (50 μm), one cell tended to pause its SS activity with saccades, while the neighboring cell tended to pause then burst. Indeed, in both cells the SSs remained modulated long after the saccade ended. However, when the target appeared to the left of the fovea, both P-cells responded with an increased probability of CS (center subplot of Fig. 1D & 1E), and when the target appeared to the right, both decreased their CS probability. Production of a CS in one P-cell was followed by 10-20 ms suppression of SS in that cell but not the neighboring cell (Supplementary Fig. 2B and 2D). Yet, the SSs shared a degree of temporal coordination: the probability of observing a SS in P-cell 2 in a 1 ms window of time increased by 39.3% if P-cell 1 happened to generate a SS during the same period (Supplementary Fig. 2C).

To quantify SS coordination during saccades, we measured the probability of synchronization with respect to chance ^6,21^. The synchronization index quantified the probability that both cells fired a spike during a 1 ms interval of time, corrected for the independent probabilities of spiking in each cell (all probabilities were conditioned on a saccade to a specific direction at time zero). Thus, the index determined whether there was greater synchrony than expected, where chance was quantified from the saccade related changes in the average firing rates of each neuron.

Remarkably, while both cells reduced their firing rates during saccades, their spikes became more synchronized (Fig. 1F). Moreover, the probability of synchronization depended on the direction of the saccade: it was greatest when the saccade was toward the direction for which complex spikes were least likely (CS+180). In these two P-cells, the probability of SS synchronization reached a maximum around the time when the saccade decelerated and came to a stop.

To analyze the data in our population, we began by measuring the CS response of each P-cell to the various visual events (primary, corrective, or central target). For each event, we estimated the target direction that produced the largest CS probability (CS-on, Fig. 2A). We found that the direction of CS-on remained consistent across the various targets (Fig. 2C, within cell comparison of direction of CS-on in response to visual event type, primary vs. secondary target 0.1°±5.4°(SEM), t(148)=0.03, p= 0.98; primary vs. central target -3.7°±5.2°, t(148)=-0.71, p= 0.48, secondary vs. central target -3.8°±5.3°, t(148)=-0.72, p= 0.47). Thus, we combined the CS response for all three visual events and used the results to define the CS-on direction of each P-cell (Fig. 1A, all targets).

**Fig. 2.**
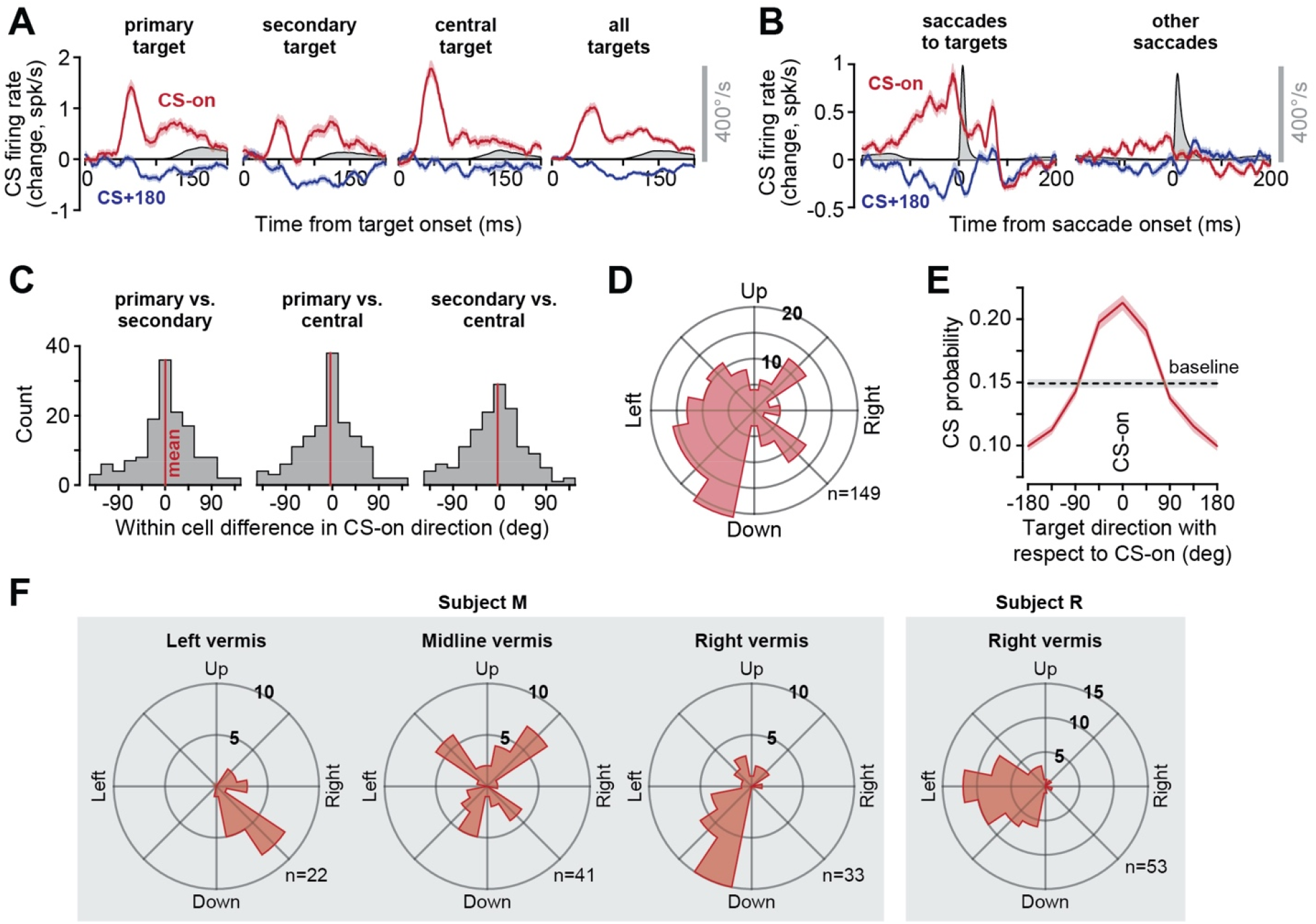
Complex spikes exhibited tuning with respect to the direction of target, and this tuning was anatomically organized. **A**. CS response aligned to target onset. For each type of target, the direction of stimulus that produced the greatest probability of CS was labeled as CS-on. Eye velocity is shown in gray. **B**. CS response aligned to saccade onset. Modulation of CS response was present before saccades that were visually instructed. The response was muted before “other saccades”. **C**. Within cell difference between CS-on directions as computed following the onset of the primary target, the secondary target, and the central target. We found no systematic differences in the estimate of CS-on between various types of targets, and thus combined the response for all targets to compute the CS-on of each P-cell. **D**. Distribution of CS-on across the population of P-cells. **E**. CS tuning function. **F**. Distribution of directions of CS-on in various regions of the vermis in two animals. Error bars are SEM.

The distribution of CS-on directions across the P-cells varied widely (Fig. 2D). However, the CS-on direction was not random. Rather, it varied with the location of the cell in the vermis: P-cells in the left vermis tended to have right-ward CS-on, while P-cells on the right had left-ward CS-on (Fig. 2F). When target directions were represented with respect to CS-on, the result was a unimodal tuning function that described the CS response of P-cells following presentation of a visual target (Fig. 2E). The probability of CS increased by 43.5±2.6% (mean±SEM) above baseline when the stimulus was presented in direction CS-on but decreased by 34.6±1.7% below baseline when it was presented in direction CS+180.

Because each target instructed a saccade, we wondered whether the CS response was due to the sudden onset of the stimulus or associated with the movement that followed. Although our experiment was not designed to specifically answer this question, we made an interesting observation. Saccades that were made in response to visual targets were preceded with large changes in CS firing rates (Fig. 2B, saccades to targets). However, saccades that were not instructed by a target, but were in the same direction and amplitude, were preceded by significantly smaller modulation of CS firing rates (Fig. 2B, “other saccades”, average CS firing rate 50ms before saccade onset in direction CS-on, paired t-test, t(296)=6.4, p=5×10^−10^, direction CS+180, t(296)=-2.8, p=0.005). Thus, the complex spikes were modulated primarily in response to sensory events that instructed movements, but not when similar movements were made spontaneously.

The simple spikes exhibited a variety of patterns during saccades: some P-cells increased their activity, some decreased their activity, while others produced more complicated patterns (Fig. 3A). The activity patterns did not separate the cells into clusters, but rather formed a continuum (Fig. 3B). For the sake of labeling, we divided the P-cells into two groups: pausers and bursters. 48% of our cells were bursters, while 52% were pausers (Supplementary Fig. 3C). To quantify how well their activities were modulated during saccades, for each P-cell we measured the change in SS rates aligned to saccade onset and computed a z-score (Supplementary Fig. 3A). Indeed, the P-cell SS rates were modulated strongly during the movements (Supplementary Fig. 3B, z-score 7.5±0.3, mean±SEM).

**Fig. 3.**
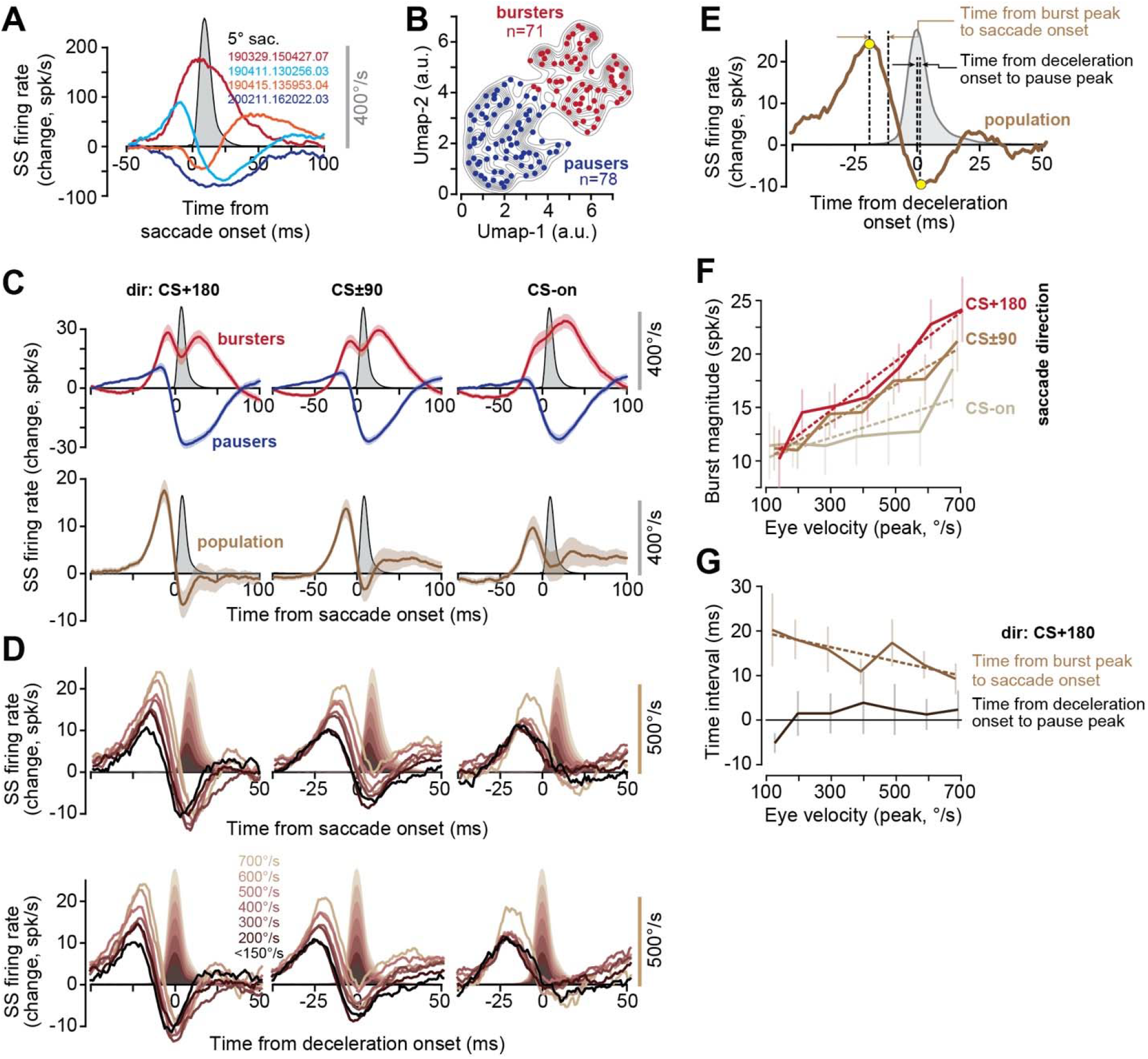
Population response of simple spikes encoded saccade direction, peak velocity, and the onset of deceleration. **A**. Average change in the firing rates of four representative P-cells with respect to baseline, during saccades (data collapsed across all directions). **B**. Clustering of saccade-aligned change in firing rates for all P-cells, using the algorithm UMAP ^51^. Separating the data into two clusters produces bursters (red) and pausers (blue). **C**. Activities of the bursters and pausers during saccade in various directions. The population response is the sum of firing rates in all P-cells. **D**. Population response aligned to saccade onset and deceleration onset. The burst tends to grow with saccade velocity and shifts forward in time, but the pause remains invariant with respect to the onset of deceleration. **E**. Quantification of the population response with respect to saccade kinematics. **F**. Magnitude of the burst before saccade onset as a function of saccade peak velocity in various directions. **G**. Timing of the burst with respect to saccade onset decreased with increased velocity, while the timing of the pause with respect to deceleration onset remained invariant.

We next organized the P-cells based on a computational model that incorporated an important anatomical feature of the cerebellum: P-cells that have similar CS tuning not only receive similar olivary inputs, but they also are likely to be part of a single olivo-cerebellar module ^22–27^. This anatomical organization implied that to estimate activity of a population of P-cells that belonged to an olivo-cerebellar module, we needed to compute saccade direction with respect to the CS-on of each P-cell. By using this coordinate transformation, we estimated the population SS response in a hypothetical olivo-cerebellar module when saccades were in direction CS-on, CS+90, etc. (Fig. 3C).

Unsurprisingly, activities of the bursters and pausers were modulated long after the saccade ended (Fig. 3C, top row). However, when the activities across all cells were organized into a population and summed, the response exhibited a clear pattern: there was a burst that preceded saccades in all directions. Notably, for direction CS+180 the burst was followed by a pause that ended near saccade termination (Fig. 3C, bottom row). This burst-pause pattern was somewhat weaker for saccades in direction CS±90, and the pause was missing entirely for saccades in direction CS-on.

The burst increased with the velocity of the impending saccade (Figs. 3D). However, the rate of increase as a function of velocity was direction dependent, showing the greatest gain for saccades in direction CS+180 (Fig. 3F, dir CS+180, F(1,5)=88.3, p=0.0002). This pattern is called a gain field, confirming earlier findings in the macaque cerebellum ^16^. Thus, before saccade onset the P-cells appeared to inhibit the nucleus with a magnitude that depended on the velocity and direction of the forthcoming saccade.

In direction CS+180, the magnitude of the burst increased with saccade velocity, but its timing shifted forward: the period from the peak of the burst to the onset of the saccade (Fig. 3E) became smaller as saccade velocity increased (Fig. 3G, r^2^=0.65, F(1,5)=9.4, p=0.027). As the saccade started toward direction CS+180, the activity changed from a burst to a pause (Fig. 3D). However, unlike the burst that preceded the saccade, the pause magnitude and timing remained invariant with respect to saccade velocity (Fig. 3G, time of deceleration onset to pause peak as a function of velocity, F(1,5)=2.4, p=0.18, Fig. 3D, rate of pause as a function of velocity, F(1,5)=1.1, p=0.34). Critically, despite a 7-fold change in velocity, the timing of the maximum pause was unchanged with respect to saccade deceleration onset (Fig. 3G). Thus, with increased saccade velocity the burst magnitude increased, and its timing shifted forward. However, regardless of saccade velocity, the pause that followed the burst was time-locked to the onset of saccade deceleration.

This invariant relationship between the timing of the pause in firing rates and the onset of saccade deceleration (in direction CS+180) raised the possibility that the P-cells were signaling when the nucleus cells should fire, presumably stopping the saccade. However, entraining the nucleus neurons would be more efficient if the P-cells not only reduced their firing rates (thus disinhibiting the nucleus), but also synchronized their spikes ^7,8^. To test this hypothesis, we computed the probability of synchronized firing in our population of simultaneously recorded P-cells.

Production of a SS in a P-cell was associated with 31% increase in the probability (with respect to chance) that there would be a simultaneous (1 ms window) SS in another P-cell (Fig. 4A, top row). Similarly, production of a CS in a P-cell increased the probability of observing a CS in another P-cell at ±5 ms latency by 227% with respect to chance (Fig. 4A, bottom row). Finally, production of a CS in one P-cell reduced the probability of SS in another P-cell at 1 ms latency by 27% (Fig. 4A, middle row). All of these observations are consistent with earlier findings in mice, demonstrating that nearby P-cells not only share a degree of spike synchrony ^6^, but that a CS in one cell can briefly suppresses SS in another cell ^28^. Furthermore, the simultaneously recorded P-cells tended to have very similar CS-on directions (Fig. 4B, between cell difference -4.4°±6.3°). However, did the P-cells synchronize their activities during saccades?

**Fig. 4.**
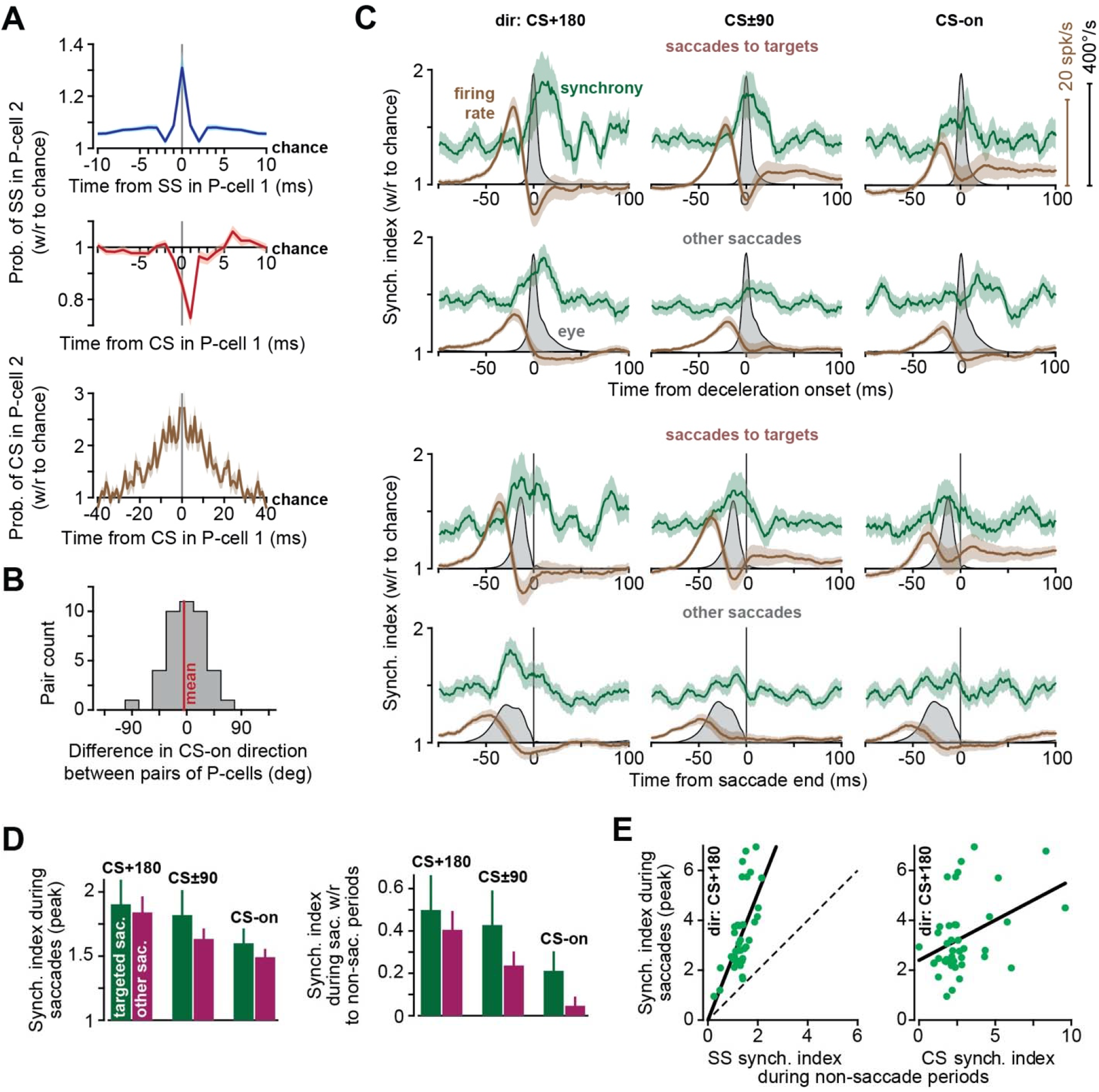
P-cells synchronize their spikes during saccade deceleration. **A**. Probabilities of spike synchronization in pairs of P-cells during the entire recording session (41±2 minutes, mean±SEM). Top: probability of simple spike in P-cell 2 at time point t (with respect to chance), given that a simple spike occurred in P-cell 1 at time zero. Middle: probability of simple spike in P-cell 2, given that a complex spike was produced in P-cell 1 at time zero. Bottom: probability of complex spike in P-cell 2 given that a complex spike was produced in P-cell 1 at time zero. Bin size is 1 ms. **B**. Difference in CS-on directions among pairs of simultaneously recorded P-cells. **C**. Synchronization index (green) and firing rates (brown) for targeted saccades and other saccades. In the top two rows, data are aligned to deceleration onset. In the bottom two rows, data are aligned to saccade end. Firing rate is the population response. Bin size is 1 ms. **D**. The magnitude of the synchronization index during saccades (peak value) for saccades in various directions. **E**. Left plot shows the magnitude of the synchronization index during saccades (peak value) in direction CS+180, with respect to SS synchronization index as measured during non-saccade periods. Dashed line is identity. Center plot shows the synchronization index during saccades with respect to complex spike synchronization index as measured during non-saccade periods (1ms bin for SS and 10ms bin for CS). Error bars are SEM.

Before saccade onset there was a burst in the P-cell population response. Surprisingly, probability of synchronization remained at baseline (Fig. 4C, note that before saccade onset, synchronization is greater than chance because at baseline the neighboring P-cells are more synchronous than chance). As the saccade started and then began to decelerate (in direction CS+180) the firing rates fell, but the synchronization index increased, reaching its peak probability after deceleration onset but before saccade end (2.3±3.5 ms after deceleration onset and -5.2±4.1 ms before saccade end). That is, during saccade deceleration the few spikes that remained were significantly more likely than chance to be synchronized. Indeed, synchronization varied with the direction of the saccade: the greatest synchronization occurred in direction CS+180 (Fig. 4D, Fig. 4D, Repeated Measure ANOVA, significant effect of direction, F(2,82)=1269.0, p=0.021). Thus, the synchronization pattern was strongest in the direction associated with the smallest probability of complex spikes.

Like targeted saccades, during other saccades the SS firing rates exhibited a burst before saccade onset, with a magnitude that was largest for direction CS+180. These saccades also had a synchronization that peaked before saccade end (Fig. 4C), with a probability that was largest for direction CS+180 (Fig. 4D). Thus, while complex spikes showed strong modulation before targeted saccades but not task-irrelevant saccades, SS firing rates were modulated during all saccades. Indeed, regardless of whether saccades were target-driven or not, simple spikes reached their greatest probability of synchrony as the movement decelerated and came to a stop.

To check whether the increased synchronization during saccades was an artifact of the change in firing rates, we performed a simulation of spiking neurons that burst and paused like cells in our population, but had independent probabilities of spike timing. The changes in rates during simulated saccades produced a synchronization index that remained at chance (Supplementary Fig. 4).

Finally, we found that pairs of P-cells that had greater CS synchrony, as measured during non-saccade periods, tended to have greater SS synchrony during saccades (Fig. 4E, F(1,40)=7.6, p=0.009). Pairs of P-cells that had greater SS synchrony during non-saccade periods also exhibited greater SS synchrony during saccades (Fig. 4E, F(1,40)=29.0, p=3×10^−6^). However, the gain of the relationship between saccade period synchrony and general synchrony was significantly greater than 1 (F(1,40)=10.7, p=0.002). Thus, although the timing of simple spikes among nearby P-cells was generally coordinated, during saccades this coordination was greatly enhanced, especially during deceleration.

## Discussion

In describing symptoms of cerebellar damage, Holmes ^29^ noted that “the most obvious errors are seen toward the end of the movement [during which] the speed of the affected limb is often unchecked until the object is reached or even passed.” For example, during an outward reach, many interposed nucleus neurons of the cerebellum produce their greatest discharge during deceleration, with spiking activity that plays a causal role in stopping the movement ^30^. Similarly, following inactivation of the fastigial nucleus, extraocular motoneurons that act as saccade agonists produce an abnormally large amount of activity during the deceleration period of ipsilateral movements ^31^, resulting in saccades that overshoot the target ^32,33^. Thus, the computations that are performed by the cerebellar cortex are critical for monitoring the ongoing motor commands and predicting when the movement should be stopped. Yet, P-cell simple spikes are often modulated long after the movement ends ^34–38^.

Here, we found that if P-cells were organized based on their complex spike response to visual stimuli, their population simple spikes produced a burst-pause pattern that started before saccade onset and ended with saccade termination. Changes in saccade velocity affected the timing and magnitude of the burst, but the pause remained time-locked to deceleration onset. Critically, in simultaneously recorded P-cells, during the pause period the probability of spike synchronization reached a maximum value. The resulting inhibition-disinhibition pattern of firing rates, coupled with spike synchronization, hints that the P-cells attempted to entrain the nucleus neurons specifically at the onset of deceleration ^7,9,39^.

What might be the behavioral consequence of this synchronization? The synchronization probability was greatest for saccades that were in the direction that coincided with the least probability of complex spikes (CS+180). For both saccades and limb movements, the CS tuning of a P-cell is likely aligned with the direction of action of the downstream nucleus neuron ^17,40^. For example, trial-to-trial analysis of the effects of complex spikes on simple spikes and behavior suggests that P-cells that have CS-on tuning to the left project to nucleus neurons that have a downstream direction of action that indirectly promotes production of leftward forces ^17^. This implies that during a saccade in direction CS+180, the increased synchrony combines with disinhibition (peak pause) to entrain the nucleus neurons during deceleration^7^, producing downstream forces that are aligned with the CS-on of the parent P-cells. As a result, the effect of synchronization of P-cells, coupled with disinhibition of the nucleus, is likely the production of forces that oppose the direction of movement, bringing it to a stop.

To our knowledge, one earlier work had reported increased spike synchrony among P-cells during movements. Using multiunit signals (i.e., not single unit isolation of spikes), Heck et al. ^10^ found increased covariance between P-cells during reaching movements (in rats). That work found that synchrony was most prominent as the hand approached the target, i.e., during deceleration. Here, we found that P-cell firing rates and spike synchrony were coordinated, especially among populations that had a common CS tuning.

Our results were obtained in the marmoset, a New World primate that like macaques and humans relies on saccades to explore its environment. Like macaques, individual P-cells in lobule VI and VII of marmosets produced simple spikes that were bursting, pausing, or a combination of the two, with no obvious relationship to the direction of the saccade or its velocity. However, following onset of a visual stimulus, the P-cells received information from the olive regarding the location of the stimulus with respect to the fovea, producing complex spikes that were tuned to the direction of the target. In both species, this tuning was anatomically organized, with P-cells in the right vermis showing highest CS probability for targets to the left. A similar anatomical representation of contralateral stimuli/movements of the arm has recently been noted in the vermis of mice ^41^.

In both marmosets and macaques, when we organized P-cells based on their complex spike tuning properties, the simple spikes produced firing rates that varied strongly with direction and velocity of the movement ^16,17^. Indeed, the gain of the response with respect to velocity was highest when saccades were in direction CS+180, and in both species the response was a burst followed by a pause that ended as the movement came to a halt. The consistency of these results across species suggests that viewing P-cell activity through the lens of population coding ^27^, i.e., a lens in which the climbing fibers organize the P-cells into olivo-cerebellar modules ^42^, may provide a key for unlocking the language with which the cerebellar cortex encodes information.

How does synchronization arise during a specific phase of a movement? P-cells that show elevated synchrony in their complex spikes also tend to fire simple spikes more synchronously ^43^. Indeed, here we found that P-cells with greater complex spike synchrony tended to have greater simple spike synchrony during saccades. In addition, we found that following a CS in one P-cell, after a 1 ms delay there was a 1-2 ms period of simple spike suppression in the neighboring P-cell, confirming recent findings in mice ^28^. One possibility is that P-cells that receive a common input from the olive generate a synchronized CS, leading to SS suppression, which may then be followed by a synchronized resumption of SS firing ^44^. However, SS synchrony was greatest in direction CS+180, i.e., the direction for which there was the smallest probability of CS. Furthermore, complex spikes showed modulation before the onset of targeted saccades but not task-irrelevant saccades, whereas simple spikes showed synchronization for both types of saccades. These observations make it seem unlikely that during saccades, presence of complex spikes played a role in synchronization of simple spikes.

Synchrony is also present in P-cells that are likely to have common parallel fiber inputs (on-beam) but different climbing fibers ^10,45^. Indeed, here we found that nearby P-cells not only had greater than chance levels of synchrony, but that cells with greater SS synchrony in general had much greater than expected SS synchrony during saccades. Thus, it is possible that SS synchrony arises from a shared input from ascending granule cell axons that are positioned directly beneath the P-cells and receive inputs from the same mossy fibers ^10^. However, why this synchrony would be focused during the deceleration phase, particularly for saccades in direction CS+180, is unclear. Because granule cells also recruit molecular layer interneurons ^46^, synchronization of P-cells may engage a network wide organization to overcome inhibition by the basket and stellate cells ^47^.

In a typical artificial neural network, information transfer from one layer to the next is via firing rates of neurons, and learning modifies synaptic weights to change the activity of each neuron and minimize error in the output layer. The cerebellum resembles a 3-layer network where learning is at least partially guided by the climbing fibers ^27,48^. Our results demonstrate that the information that is transmitted from the P-cells to the nucleus is encoded in an exquisite coordination of firing rates and synchronization. This implies that when there is error in performance, cerebellar learning cannot simply focus on changing the P-cell firing rates. Rather, learning must also alter network wide synchronization. This conjecture predicts that complex spikes that arise following movement errors not only promote learning via changes in the activity of individual P-cells ^17,49,50^, but may also alter the synchronization patterns of populations of P-cells. Learning to transfer information via synchronization is an exciting new direction with which to explore the function of the cerebellum.

## Acknowledgements

The work was supported by grants from the National Science Foundation (CNS-1714623), the NIH (R01-EB028156, R01-NS078311), and the Office of Naval Research (N00014-15-1-2312).

## Author contributions

E.S.N., J.S.P., P.H., and R.S. conceived and performed experiments. E.S.N., J.S.P., M.A.F., and R.S. analyzed data, E.S.N. made figures, performed statistical analysis, and performed simulations, R.S. and E.S.N. wrote the manuscript.

---

**Supplementary Fig. S1.**
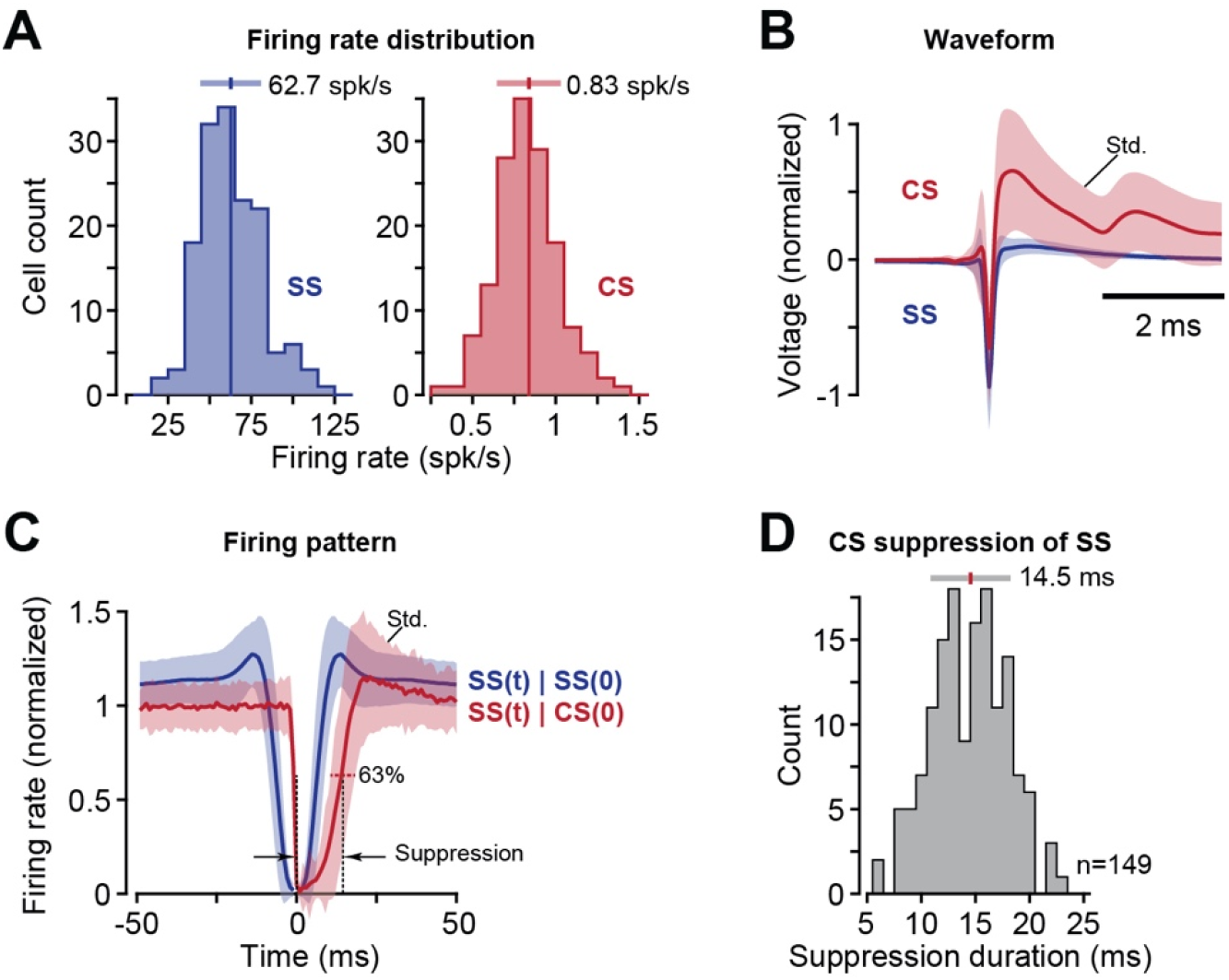
Properties of saccade-related P-cells (n=149) in the marmoset vermis lobule VIa and VIIa-c. **A**. Distribution of average firing rates for simple spikes (blue) and complex spikes (right). The bar at the top indicates mean and standard deviation. **B**. Waveforms for the simple and complex spikes. The waveforms were normalized by setting the cell’s mean voltage to 0 and the maximum negative going simple spike deflection to -1. Error bars are standard deviation. **C**. Within-cell interactions between simple and complex spikes. The blue curve shows the firing rate of simple spikes at time t, given that the cell produced a simple spike at time zero, labeled as SS(t)|SS(0). The red curve shows the firing rate of simple spikes at time t, given that the cell produced a complex spike at time zero, labeled as SS(t)|CS(0). Simple spike rates for each P-cell were normalized with respect to average simple spike firing rate as computed over the entire recording session. Error bars are standard deviation. **D**. Suppression period of simple spikes following production of a complex spike. Suppression period for each P-cell was defined as the duration of time after a complex spike that was required before the simple spike firing rate recovered 63% of its pre-complex spike value. The bar at the top indicates mean and standard deviation.

**Supplementary Fig. S2.**
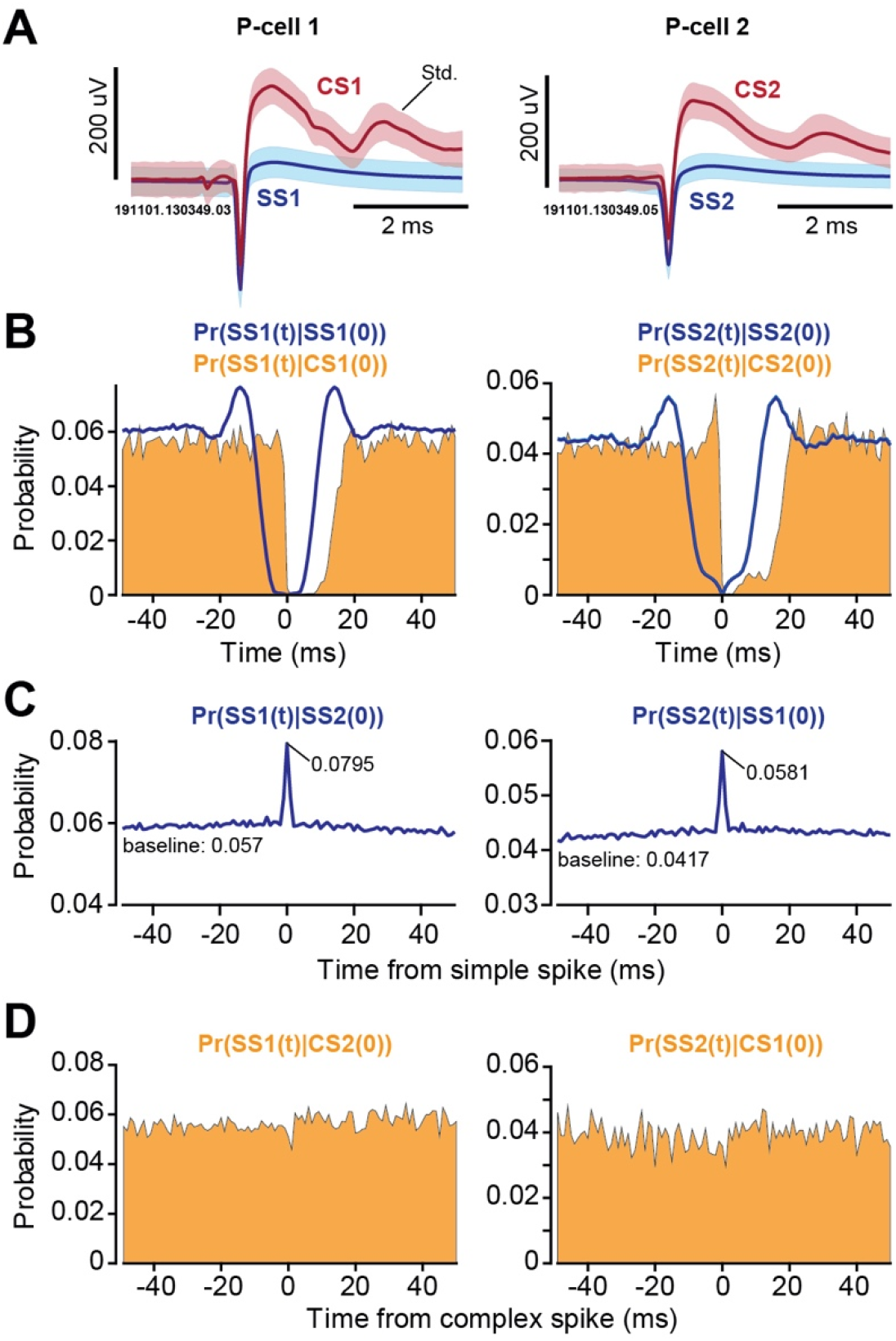
Spike timing properties of a sample pair of simultaneously recorded P-cells. These are the same cells as in Fig. 1. **A**. Simple and complex spike waveforms. Error bars are standard deviation. **B**. The curve Pr(SS1(t)|SS1(0)) quantifies the probability of a simple spike in P-cell 1 at time t, given that P-cell 1 produced a simple spike at time zero. This quantifies the simple spike refractory period. The curve Pr(SS1(t)|CS1(0)) quantifies the probability of production of a simple spike in P-cell 1 at time t, given that P-cell 1 produced a complex spike at time zero. This indicates the complex spikes induced suppression of simple spikes. **C**. The curve Pr(SS1(t)|SS2(0)) quantifies the probability of production of a simple spike in P-cell 1 at time t, given that a simple spike was produced by P-cells 2 at time zero. **D**. The curve Pr(SS1(t)|CS2(0)) quantifies the probability of production of a simple spike in P-cell 1 at time t, given that a complex spike was produced by P-cells 2 at time zero. Bin size is 1 ms in the probability plots.

**Supplementary Fig. S3.**
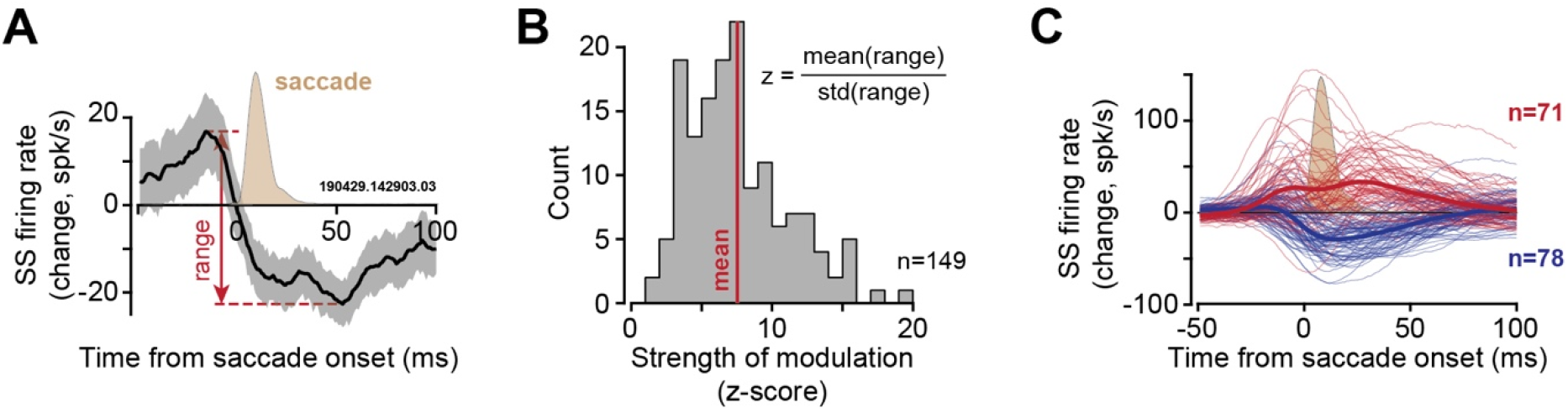
Modulation of simple spikes in individual P-cells during saccades. **A**. Data from a single P-cell, quantifying the change in simple spike rates, aligned to saccade onset. Range was defined as the maximum change in firing rate in the pre-to post-saccade period. The light brown curve shows the average saccade velocity (peak value is 475 deg/s). Error bars are standard deviation computed via bootstrapping. **B**. Strength of saccade-related modulation of each P-cell was defined via a z-score. The z-score computed the mean of the range of the firing rates divided by the standard deviation of the range (7.5±0.3 mean±SEM). Strongly modulated P-cells were those that had a z-score greater than 3, composing 96% (142 out of 149) of the cells in the population. **C**. Change in simple spike rates with respect to baseline for the bursters (red) and pausers (blue). Baseline firing rate is defined as the average firing rate as measured during the entire recording session.

**Supplementary Fig. S4.**
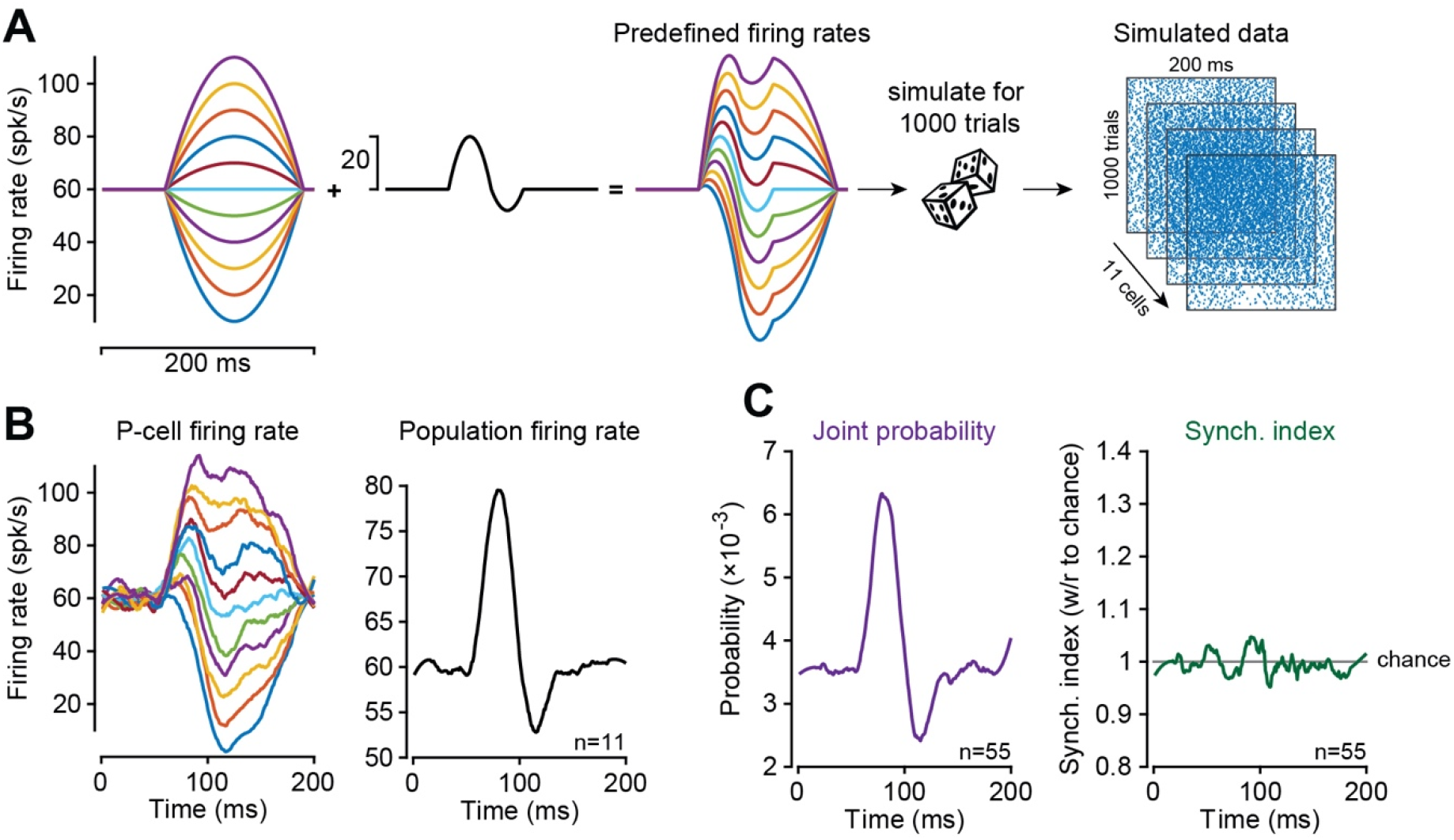
To check whether the increased synchronization index during saccades was an artifact of the change in firing rates of P-cells, we performed a simulation of spiking neurons that burst and paused like the cells in our population. However, the simulated cells had independent probabilities of spiking. **A**. The simulated population consisted of 11 neurons ranging from bursting to pausing in their activity. The plots show the construction of the average firing rates. We started with a unimodal function of various amplitudes, added the response in the middle row to each neuron, producing the data on the right, representing the instantaneous average firing rate 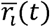 of each simulated neuron. **B**. Data from 1000 simulated trials. The plot shows the average firing rate of each simulated cell. **C**. The population firing rate of the simulation. **C**. Joint probability, and synchrony index of 55 pairs of cells (11 cells taken 2 at a time). Joint probability reflected modulation of firing rates, but because spike timing in each neuron was independent, the synchrony index remained at chance level.

## Methods

Neurophysiological data were collected from two marmosets (*Callithrix Jacchus*, male and female, 350-370 g, subjects M and R, 6 years old. The marmosets were born and raised in a colony that Prof. Xiaoqin Wang has maintained at the Johns Hopkins School of Medicine since 1996. The procedures on the marmosets were evaluated and approved by the Johns Hopkins University Animal Care and Use Committee in compliance with the guidelines of the United States National Institutes of Health.

### Data acquisition

Following recovery from head-post implantation surgery, the animals were trained to make saccades to visual targets and rewarded with a mixture of applesauce and lab diet ^18^. Visual targets were presented on an LCD screen (Curved MSI 32” 144 Hz - model AG32CQ) while binocular eye movements were tracked using an EyeLink-1000 eye tracking system (SR Research, USA). Timing of target presentations on the video screen was measured using a photo diode.

We used the MRI and CT imaging data for each animal and designed an alignment system that defined trajectories from the burr hole to various locations in the cerebellar vermis, including points in lobule VI and VII. We used a piezoelectric, high precision microdrive (0.5 micron resolution) with an integrated absolute encoder (M3-LA-3.4-15 Linear smart stage, New Scale Technologies) to advance the electrode.

We recorded from the cerebellum using four types of electrodes: quartz insulated 4 fiber (tetrode) or 7 fiber (heptode) metal core (platinum/tungsten 95/05) electrodes (Thomas Recording), and 64 channel checkerboard or linear high density silicon probes (M1 and M2 probes, Cambridge Neurotech). We connected each electrode to a 32 or 64 channel head stage amplifier and digitizer (RHD2132 and RHD2164, Intan Technologies, USA), and then connected the head stage to a communication system (RHD2000 Evaluation Board, Intan Technologies, USA). Data were sampled at 30 kHz and band-pass filtered (2.5 - 7.6 kHz). We used OpenEphys ^52^, an open-source extracellular electrophysiology data acquisition software, for interfacing with the RHD2000 system and recording of signals.

### Behavioral protocol

Each trial began with fixation of a center target for 200 ms, after which a primary target (0.5×0.5 deg square) appeared at one of 8 randomly selected directions at a distance of 6.5 deg. Onset of the primary target coincided with presentation of a distinct tone. As the animal made a saccade to this primary target, that target was erased, and a secondary target was presented at a distance of 2-2.5 deg, also at one of 8 randomly selected directions. The subject was rewarded if following the primary saccade it made a corrective saccade to the secondary target, landed within 1.5 deg radius of the target center, and maintained fixation for at least 200 ms. Onset of reward coincided with presentation of another distinct tone. Following an additional 150-250 ms period (uniform random distribution), the secondary target was erased, and the center target was displayed.

### Data analysis

All saccades, regardless of whether they were instructed by presentation of a visual target or not, were identified in the behavioral data using a velocity threshold. Saccades to primary, secondary, and central targets were labeled as targeted saccades, while all other saccades were labeled as task irrelevant.

Electrophysiological data were sorted into spikes using P-sort ^53^, a newly developed open-source software that we developed specifically for identification of simple and complex spikes. P-sort provides tool to help confirm that the complex and simple spikes originate from the same P-cell. Briefly, we compared the conditional probability *Pr* (*S*(*t*)|*C*(0)) with *Pr* (*S*(*t*)|*S*(0)). For example, *Pr* (*S*(*t*)|*C*(0)) is the probability that a simple spike occurred at time *t*, given that a complex spike was generated at time zero. *Pr* (*S*(*t*)|*S*(0)) is the probability that a simple spike occurred at time *t*, given that a simple spike was generated at time zero. Simple spikes that originate from a single P-cell produce a refractory period. Thus, *Pr* (*S*(*t*)|*S*(0)) exhibits a low probability period of roughly 5 ms in duration after time zero. On the other hand, a complex spike coincides with suppression of future simple spikes, but not those that occurred before. As a result, *Pr* (*S*(*t*)|*C*(0)) is asymmetric, with a long period of low simple spike probability (around 15 ms) following time point zero. Examples are provided in Supplementary Fig. S2.

Simple and complex spike baseline firing rates were computed by dividing the total number of spikes by the duration of the entire recording. Simple and complex spike instantaneous firing rate were calculated from peri-event time histograms with 1 ms bin size. Events of interest included: visual events (target onset), saccade onset, deceleration onset, and saccade offset. We used a Savitzky–Golay filter (2nd order, 31 datapoints) to smooth the traces for visualization purposes.

Complex spike tuning was computed by measuring the CS probability following target onset. We counted the number of complex spikes after target onset up to saccade onset or a fixed 200 ms window, whichever happens first. This approach ensured that the complex spikes during saccades or after saccade offset did not get included in the measurements. Dividing the spike count by the number of events resulted in the CS probability in each direction.

Suppression duration of simple spikes following a complex spike (Supplementary Fig. S1C) was computed by measuring the period during which the simple spikes recovered 63% of their pre-complex spike firing rate.

To compute population response during saccades, we began by computing the change in simple spike firing rate of each P-cell with respect to its baseline. Next, we labeled each saccade by measuring its direction with respect to the CS-on of the recorded P-cell. Finally, we summed the activities in all P-cells (i.e., changes with respect to baseline) for saccades in direction CS-on, CS+45, etc., using a bin size of ±25 deg.

### Analysis of the simultaneously recorded P-cells

Multi-channel electrodes allowed for analysis of simultaneously recorded neurons. However, spiking activity in one neuron can easily influence the data recorded by two nearby channels, thus giving an illusion that the two channels are picking up two distinct neurons. To guard against this, after we sorted the data in each channel, we waveform triggered the data recorded by channel A by the spikes recorded on channel B. This identified the waveform of the neuron recorded by channel B on the spike recorded on channel A. We compared this cross-channel triggered waveform with the within channel triggered waveform generated by the spikes recorded by channel A. The cross-channel triggered waveform must produce a different cluster of spikes in A than the main neuron isolated by A. If there were spikes that occurred within 1 ms of each other on channels A and B, we used these coincident-spike events to trigger the waveform in A. The spikes in A that were identified to be coincident with B should look approximately the same as the non-coincident spikes in A. Examples of this approach are provided in Sedaghat-Nejad et al. ^18^.

To quantify coordination between activities of two P-cells, we computed joint probabilities, corrected for chance ^6,21^. We computed Pr(*S*2(*t*), *S*1(0)) */*(Pr(*S*2) Pr(*S*1)), which is equal to Pr(*S*2(*t*)|*S*1(0))*/* (Pr(*S*2)). This quantified whether the occurrence of a simple spike on channel 1 at time zero altered the probability of simple spikes on channel 2 at time t, corrected for probabilities expected from their average firing rates. Because channel labels 1 or 2 are interchangeable, we considered the average of the two cases as the corrected conditional probability for a pair of P-cells. We implemented a similar analysis to quantify the coordination between complex spikes in two cells, or complex spikes in one cell and simple spikes in another cell.

To compute the probability of synchronization of simple spikes during saccades, we began by computing the joint probability of spiking at time t, given that a saccade took place at time zero in a particular direction, Pr(*S*1(*t*), *S*2(*t*)|*sac*(0)). To correct for the fact that firing rates changed during the saccade, we divided the joint probability by the independent probabilities of spike production in each cell, measured when a saccade took place in the given direction at time zero. Thus, the synchronization index was defined for each saccade direction as:

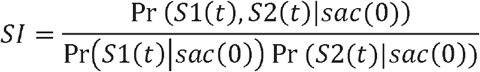

### Modeling

To check whether the increased synchronization index during saccades was an artifact of the change in firing rates of P-cells, we performed a simulation of spiking neurons that burst and paused like cells in our population but had independent probabilities of spike timing. We pre-defined the firing rate pattern for 11 hypothetical neurons all with a 60 spk/s baseline firing rate (Supplementary Fig. S4A) and then add a 130 ms duration modulation. The result produced a population response firing pattern that mimics the P-cells in our dataset (Supplementary Fig. S3C). Using a Bernoulli process and pre-defined firing rates, we simulated the spiking activity for the 11 hypothetical neurons for 1000 trials. Next, we used the methods described above to compute the estimated firing rate of each cell and the population response (Supplementary Fig. S4B). Finally, we computed the joint probability as well as the synchronization index between 55 (2 choose 11) pairs of cells (Supplementary Fig. S4C). Our results confirmed that while the joint probability was modulated according to the change in firing rates in the population of cells, the synchronization index stayed at chance level.

